# Winter agricultural practices matter : rice stubble cover and landscape feature diversity increase abundance of wintering farmland birds in the Camargue

**DOI:** 10.64898/2026.11.21.768284

**Authors:** Pierre Mallet, Thomas Galewski, Clélia Sirami, François Mesléard, Thomas Blanchon, Elie Gaget, Arnaud Béchet

## Abstract

Agricultural intensification has led to a decline of farmland bird populations worldwide. Most studies have focused on the role of practices on birds during the breeding season but little is known about the role of practices during winter. Here, we study the effect of different types of winter field cover and landscape features on wintering birds in the rice production area of the Camargue (Rhône delta, France). We conducted bird surveys in 86 organic crop fields during winter. We hypothesized that the type of field cover (rice stubble, bare ground, winter wheat, grasslands), and the type of landscape features (hedegrows, grass strips and reed strips) have an effect on the abundance and species richness of wintering bird communities at field level. Moreover, we hypothesized that the diversity of field cover types and landscape features at the landscape level have a positive effect on the abundance and species richness of wintering bird communities, in line with the landscape complementation hypothesis. Results show that at the field level, rice stubble fields accommodate higher bird abundance that fields with other cover types. Moreover, we show that this pattern is driven by a few gregarious and granivorous wintering species while others preferred grasslands. At the landscape level, we show that the diversity of landscape features increased wintering bird abundance. We also show species-specific responses to different types of landscape features with hedgerows, reed strips and grass strips affecting diverse species at both field and landscape levels. Overall, our results suggest that rice stubble provide important winter habitats for declining granivorous species, while maintaining a heterogeneous mosaic of landscape features benefits wintering bird communities. Finally, we quantify species-specific response to the surface of different landscape features, a key step to build-up dedicated agri-environmental schemes.

## Introduction

Agricultural intensification has led to a decline of farmland bird populations worldwide (IUCN 2024; PECBMS 2025). Heavy pesticide use, increased mechanization and landscape homogenization have been shown to have negative effects on breeding birds (Newton 2017; Moreau et al. 2022), while the impact of agricultural intensification on birds over the non-breeding season has received comparatively less attention (Wilson et al. 1996; Geiger et al. 2010; Katayama et al. 2015).

The large-scale switch from spring-sown to autumn-sown crops that occured in the 1970s is often viewed as a key driver of the decline of farmland birds presumably through food shortage (Chamberlain et al. 1999; Benton et al. 2002; Whittingham and Evans 2004), which decreased their winter survival (Donald and Evans 1994; Wilson et al. 1996; Peach et al. 1999; Siriwardena et al. 2008a). This might be especially true for seed-eating species, which partly rely on post-harvest stubble in winter, including several species that have declined sharply in Europe in recent decades as Eurasian tree sparrow (*Passer montanus*), Corn bunting (*Emberiza calandra*) or Yellowhammer (*Emberiza citrinella*; (PECBMS 2025)).

Landscape homogenization also seems to have influenced the composition of bird communities wintering in farmland (Geiger et al. 2010). This likely results from the negative effect of landscape homogenization on habitat complementation processes, i.e. the fact that individuals have access to complementary resources when the landscapes include a diversity of land cover types (Dunning et al. 1992; Morales et al. 2015). Indeed, agricultural intensification has led to an increase in field size, a decrease in crop diversity and a decrease in the proportion of semi-natural cover. In the dry cereal plain of Central Spain species richness and abundance of wintering birds are positively influenced by field cover diversity (e.g. bare ground, stubble, winter crops…) and the proportion of semi-natural cover in the landscape (Morales et al. 2015).

Previous studies testing the habitat complementation hypothesis during winter did not explicitly address the role of landscape features for wintering birds. Despite their relatively small proportion in the landscape, landscape features represent important habitat for wintering farmland birds. In Poland, landscape features account for 6.6% of the area of arable landscapes but support 35.6% of the abundance of wintering farmland birds (Wuczyński and Wuczyński 2019). Landscape features are used as an additional food resources area because they are usually not sprayed with synthetic inputs and are usually a less disturbed habitat, resulting in high seed densities (Vickery et al. 2002; Geiger et al. 2010).

Studies on the effect of wintering agricultural practices, in particular field cover types and landscape features, on farmland wintering birds in regions of dry cereal production are scarce (Henderson et al. 2004; Baker et al. 2012; Morales et al. 2015), but the question has received even less attention in rice production areas (Rodier et al. 2025). Rice is cultivated either in monoculture or in rotation over 5 to 7 years with wheat, alfalfa and vegetable crops (Rodier et al. 2025). Depending on rotation stage and on farm-specific management strategies, winter field cover may be stubble rice, bare ground ready for a new spring crop, grassland generally resulting from 2-3 years of alfalfa production or freshly sown winter wheat.

Landscape features may be much more diverse in rice production landscapes, with the frequent presence of hedgerows, grass strips and reed strips between fields due to the network of irrigation and drainage ditches. Both reed strips and hedgerows can be used as shelter against predators (Haviland et al. 2016) while grass strips may provide resources for insectivorous and granivorous birds (Vickery et al. 2009; Mallet et al. 2023). Landscape features may provide complementary resources, with crop fields primarily providing feeding opportunities while landscape features provide additional food resources, shelter and refuge from predators. Hence, landscape features may strongly contribute to habitat complementation processes in rice production landscapes. Thus, it is important to assess which combinations of field cover types and landscape feature types are most effective for the conservation of wintering farmland birds, to set up effective agri-environmental schemes in rice production landscapes.

In this paper, we study the effect of winter field cover types and landscape features on wintering birds in the rice production area of the Camargue (Rhône delta, France). This region hosts 103 species of wintering passerines, including high numbers of Meadow pipit *Anthus pratensis* and Corn bunting *Emberiza calandra* (Issa and Muller 2015; Fraixedas et al. 2019). We conducted bird surveys in 86 organic crop fields during winter. We hypothesized that the type of field cover (rice stubble, bare ground, winter wheat, or grassland), and the type of landscape feature (hedgerows, grass strips, reed strips) at both field and landscape levels have an effect on the abundance and species richness of wintering bird communities. We predicted that (i) bird abundance is higher in fields with rice stubble due to the expected higher availability of seeds in this field cover type; (ii) species richness is higher in landscapes with more diverse field cover types; (iii) species abundance and richness is higher in landscapes with more diverse landscape features due to landscape complementation processes; and (v) response to field cover types, landscape features and their diversity at the landscape level vary among species.

## Material and methods

### Study area and sampled field selection

Our study was conducted in the Rhône River delta, a 180,000 ha polderized flood plain located in Southern France. As part of the Natura 2000 network, the Camargue is recognized under both the Habitats and Birds Directives as Site of Community Importance and Special Area of Conservation, respectively. The area is also a Ramsar site, a Man and Biosphere reserve and a Natural Regional Park of 100 000 ha composed of 56 000 ha of semi-natural area and 46 000 ha of agricultural area (PNRC 2016). In 2020-21 rice accounted for 13 000 ha, with 21% (2750 ha) cultivated under the organic farming label. Rice is sown in May and flooded from May to October. Rice cultivation allows washing out salt-rich soils and allows rotation with dry crops, mainly wheat and alfalfa. The classic organic crop rotation in Camargue is rice, followed by winter wheat and alfalfa (Mouret 2018). Spring-sown crops such as sunflower and vegetables may also be included in the rotation. Winter wheat is sown from October to January. Winter wheat does not systematically follow rice harvest so that rice stubble may last until spring, when soil preparation for the next spring crop occurs. Alfalfa is generally sown in fall and remain for two to three years with several annual cuts.

Landscape features are an important component of agricultural landscapes in Camargue. In order to retain water into crop fields, fields of rice paddy landscapes are often separated by wide (> 3 m) reed strips, hedgerows or grass strips often co-occur (Mallet et al. 2023).

We selected only organic crop fields to limit potential biases related to variations in the intensity of agricultural practices (Fig. 1). To select sampled crop fields, we built a land cover map of the study area using 2019 data from the BD TOPO^®^, the OSO Land Cover Map and the Camargue Regional Natural Park. We estimated the area of semi-natural cover and the length of hedgerows (the only type of landscape feature for which a GIS layer was available) in a 500m^2^ moving window with a step size of 100 m. We then used the method developed by (Pasher et al. 2013) to select a combination of 86 crop fields that maximize the gradients of semi-natural cover and hedgerow length, while minimizing the correlation between them.

**Figure 1.**
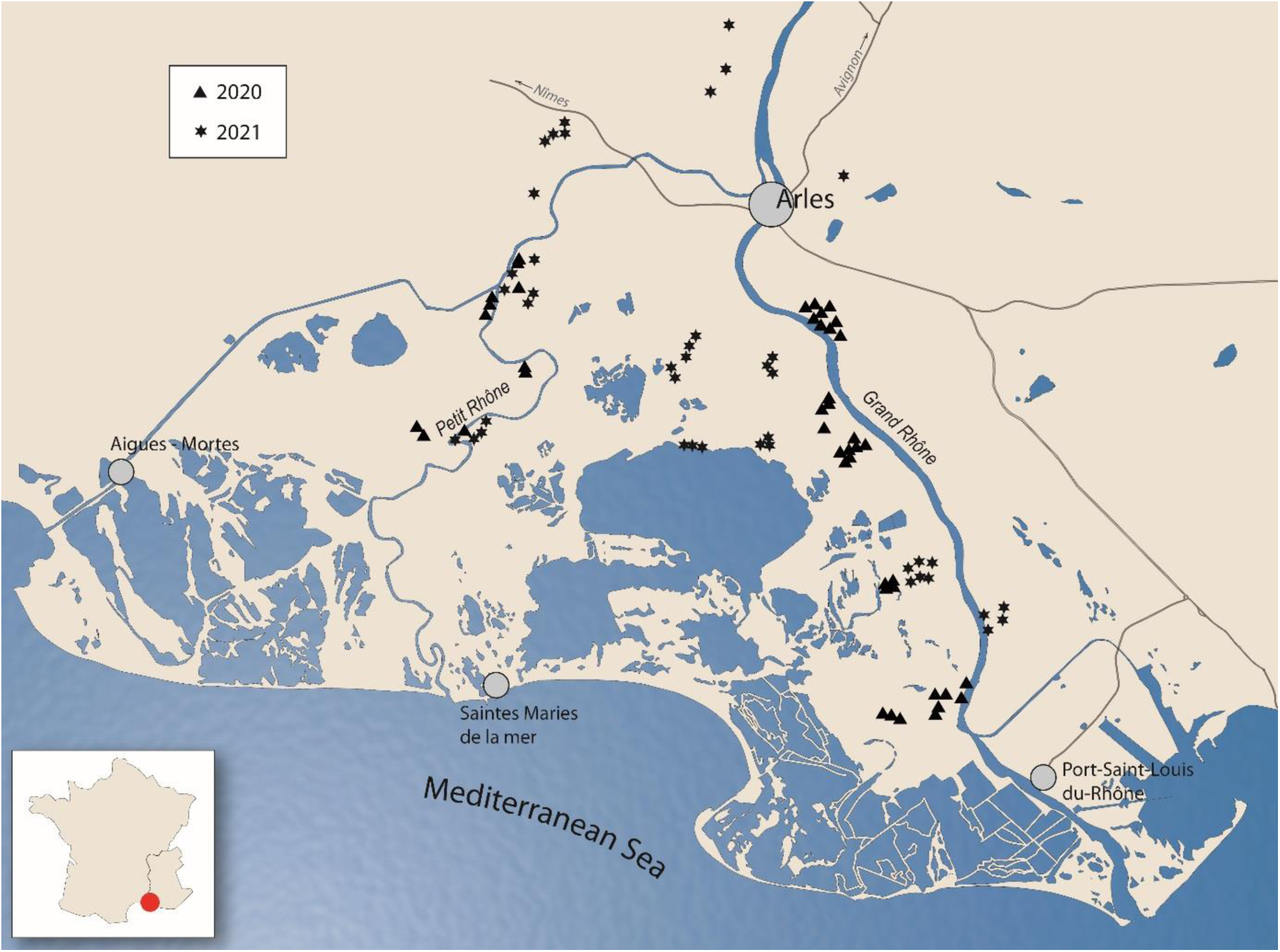
Location of the eighty-six fields selected for winter bird survey in the Camargue, Rhône delta. Triangles represent crop fields monitored in the 2020-21 winter and stars crop fields monitored in the 2021-22 winter.

### Field cover types and landscape features

We mapped winter field cover during bird surveys (n=172), i.e. once between mid-November and mid-December and once in January since field cover may change along winter time (stubbles are ploughed, winter crops are sowed…). We mapped semi-natural cover types and landscape features by direct observation in the field in June 2020 and June 2021.

Winter field cover were classified into four categories 1) rice stubble (n = 29); 2) bare ground when the crop field was ploughed (n = 51); 3) winter wheat for crop fields with freshly sown, sparse winter cereals 5-10 cm high (n = 34); and 4) grassland for alfalfa and other sown hayfield with vegetation between 5 and 20 cm height (n = 58). A fifth category was used only for crop fields within the landscape: 5) woody crops, consisting of vineyards and orchards.

We characterized landscape features at the field level by mapping the surface of hedgerows, grass strips and reed strips in a buffer of 10 meter along the edge of sampled crop fields. Following Morales et al. (2015), we also measured the surface area of landscape features within a 500 m radius buffer centered on each sampled crop field.

### Winter bird surveys

Birds were counted on 86 fields in winter 2020-21 (n=45) and 2021-22 (n=41) during 10-minute walking transects along the edges of each sampled crop field. Wintering passerines were detected by calls, or observed on land while eating or in flight when flushed by the observer. The area covered during the surveys was similar across all fields and we did not find any correlation between bird abundance and crop field size (Pearson correlation coefficient < 0.19). Two visits were conducted in each sampled crop field between mid-November and January with at least 2 weeks between the two visits, following the French common wintering bird census scheme (Jiguet 2014). The same professional ornithologist (T. Blanchon) conducted all surveys. Birds flying high over the area were excluded from further analyses because they were not interacting directly with the landscape around the field. Observations of birds outside the field or beyond the field margin were also discarded to avoid detection bias potentially generated by hedgerows preventing the observer from seeing birds beyond trees.

## Data analysis

### Effects of field cover types and landscape features on abundance and species richness of wintering birds

We used generalized linear mixed models (GLMM) with total abondance and species richness calculated per field visit on each crop as response variables. We included field cover type, and, at the landscape level, the total surface of stubble area, the surface of semi-natural areas, field cover diversity, and landscape feature diversity as fixed effects (Table 1). Random effect was visit number nested into field identity. To model total bird abundance (over all species) we used a negative binomial distribution of the error term while we used a Poisson distribution for the error term when modelling species richness.

**Table 1.** Description of the explanatory variables at field and landscape level used for the linear mixed-effect modeling.

| Variable | Description | Level | Median [range] |
|---|---|---|---|
| Field level |  |  |  |
| Field cover type | Factor with four levels (rice stubble, bare ground, winter wheat, grassland) | Field | - |
| Hedgerow10 (m<sup>2</sup>) | Hedgerow area around the field, i.e. area covered by tree lines and bushy areas | 10 m | 790<br>[0 ; 7 017] |
| Reed_strip10 (m<sup>2</sup>) | Reed strips area around the field, i.e. area covered by reeds growing in and along irrigation or drainage earthen ditches | 10 m | 2 285<br>[0 ; 9 283 ] |
| Grass_strip10 (m<sup>2</sup>) | Grass strips area around the field. |  | 1 319<br>[0 ; 7 987] |
| Landscape level |  |  |  |
| Hedgerow500 (m<sup>2</sup>) | Hedgerow area in the landscape, i.e. area covered by tree lines and bushy areas | 500 m | 32 295<br>[0 ; 109 542] |
| Reed_strip500 (m<sup>2</sup>) | Reed strip area in the landscape, i.e. area covered by reeds growing in and along irrigation or drainage earthen ditches | 500 m | 30 348<br>[0 ; 73 166] |
| Grass_strip500 (m<sup>2</sup>) | Grass strip area in the landscape, i.e. area covered by grass | 500 m | 30 679<br>[0 ; 62 676] |
| Stubble area (m<sup>2</sup>) | Area of stubble fields in the landscape | 500 m | 50 789<br>[0 ; 412 315] |
| Semi-natural area (m<sup>2</sup>) | Area of all types of semi-natural cover types (grassland, woodland, wetland) in the landscape | 500 m | 142 613<br>[0; 544 033] |
| Landscape feature diversity | Shannon diversity index: - $\sum_{i=1}^{n} pi \ln pi$, where *pi* is the proportion of landscape feature *i* (hedgerow, grass strip and reed strip) in the landscape | 500 m | 0.93<br>[0; 1.09] |
| Field cover diversity | Shannon diversity index: - $\sum_{i=1}^{n} pi \ln pi$, where *pi* is the proportion of field cover type *i* in the landscape | 500 m | 0.99<br>[0.1; 1.6] |

### Species specific response to field cover types and landscape features

We assessed how each species responded to field cover types and landscape features by building a GLMM with the response variable being bird abundance per species, per field and per visit. To avoid extreme cases of zero-inflation, we only kept species present in more than 15 % of the sampled crop fields (Marja and Herzon 2012). The resulting dataset contain 13 species (Eurasian skylark *Alauda arvensis*, Meadow Pipit *Anthus pratensis*, Common buzzard *Buteo buteo*, Cetti’s warbler *Cettia cetti*, Corn bunting *Emberiza calandra*, Reed bunting *Emberiza schoeniclus*, European robin *Erithacus rubecula*, Common chaffinch *Fringilla coelebs*, Great tit *Parus major*, House sparrow *Passer domesticus*, Common chiffchaff *Phylloscopus collybita*, European stonechat *Saxicola rubicola*, and Common starling *Sturnus vulgaris*).

We considered a model with field cover explanatory variables at two levels : field cover type in the sampled field and stubble area in the landscape because they were the effects retained in the first analysis (see 2.4.1 and results in 3.1). We added an interaction term between these two explanatory variables and species identity. We also added hedgerows, grass or reed strips at both the field (measured along the 10 m buffer around the field) and landscape level (measured within the 500m buffer). Because the effect of landscape features may vary depending on species, we added an interaction term between species identity and each of these latter explanatory variable (see Table 1).

Explanatory variables were poorly correlated with each other (Pearson correlation coefficients < 0.5), except the area of reed strip and grass strip, which were positively correlated between the field level and the landscape level (r = 0.54 and 0.64 respectively, Appendix A). We used a negative binomial error distribution (type 2: variance increases quadratically with the mean) to deal with the overdispersion of abundances and a Poisson distribution for species richness. In all models, we used visit number nested into field identity as random effect. We accounted for spatial autocorrelation by using an exponential structure on crop field coordinates (Kristensen and McGillycuddy 2022), and checked for the absence of autocorrelation in the residuals. All explanatory variables (Table 1) were centered and scaled prior to model selection. We ran models with all combinations of effects which could be derived from a general model and ranked models using the Akaike information criterion AICc (Burnham and Anderson 2002). Then, we averaged parameters of models with a ΔAICc smaller than two. We discussed the effect of variables when the confidence intervals of their full averaged estimate did not include zero. We run post-hoc tests to compare species-specific abundance variations depending on field cover type. Landscape feature effects at the field and landscape level were assessed by the species-specific slope of the effects. To assess size effect and how much surface area of different landscape feature types increase species abundance, we run again the best model with unscaled variables.

Statistical analyses were run using glmmTMB (Magnusson et al. 2020), entropart (Marcon and Hérault 2015), MuMin (Barton 2020) and emmeans (Lenth et al. 2022) in R 4.3.2 (R Core Team 2017).

## Results

A total of 65 bird species were detected and recorded over the wintering surveys. Total abundance per field and per visit was on average 47.5 ± 5.4, range [0 ; 416]. Species richness per field and per visit was on average 5.9 ± 0.2, range [0; 14]. The most frequent species across fields were passerines species (see Electronic Supplementary Material ESM).

### Effect of field cover type at the field and landscape level

Model averaging suggests that total bird abundance is explained by field cover type and increases with stubble area in the landscape (β = 0.40 ± 0.14) and landscape feature diversity (β = 0.19 ± 0.13; Table 1). At field level, stubble cover supported the highest total bird abundance and up to 2.8 higher bird abundance than winter wheat (*p* = 0.02) while other pairwise comparisons were not significant (Fig. 2a). Field cover diversity and the area of semi-natural habitat were not retained by model selection (confidence interval including zero for Field cover diversity - ESM Table S2 - and ΔAIC >2 for models including semi-natural area; Table 1).

**Table 1.** Model name, fixed effects, number of parameters (k), AICc and ΔAICc for models assessing the effects of field cover type and landscape features on overall wintering bird abundance (negative binomial distribution) and community richness (Poisson distribution). Only models within 3 AICc points of the best model are presented.

| Model | Fixed effects | k | AICc | ΔAIC |
|---|---|---|---|---|
|  | **Total abundance** |  |  |  |
| **M1** | Landscape feature diversity + Stubble area | 8 | 1624.8 | 0 |
| **M2** | Field cover type + Landscape feature diversity + Stubble area | 11 | 1625.6 | 0.74 |
| **M3** | Field cover type + Field cover diversity + Stubble area | 11 | 1626.6 | 1.73 |
| **M4** | Field cover type + Field cover diversity + Landscape feature diversity + Stubble area | 12 | 1626.6 | 1.75 |
| **M5** | Field cover diversity + Landscape feature diversity + Stubble area | 9 | 1626.7 | 1.83 |
| **M6** | Semi-natural area + Landscape feature diversity | 9 | 1627.0 | 2.23 |
| **M7** | Field cover type + Stubble area | 10 | 1627.4 | 2.56 |
| **M8** | Field cover type + Landscape feature diversity + Semi-natural area + Stubble area | 12 | 1627.7 | 2.88 |
|  | **Species richness** |  |  |  |
| **M1** | Field cover type + Field cover diversity | 9 | 834.5 | 0 |
| **M2** | Field cover type | 8 | 834.9 | 0.48 |
| **M3** | Field cover type + Landscape feature diversity | 9 | 835.9 | 1.48 |
| **M4** | Field cover type + Field cover diversity + Stubble area | 10 | 836.7 | 2.21 |

**Figure 2.**
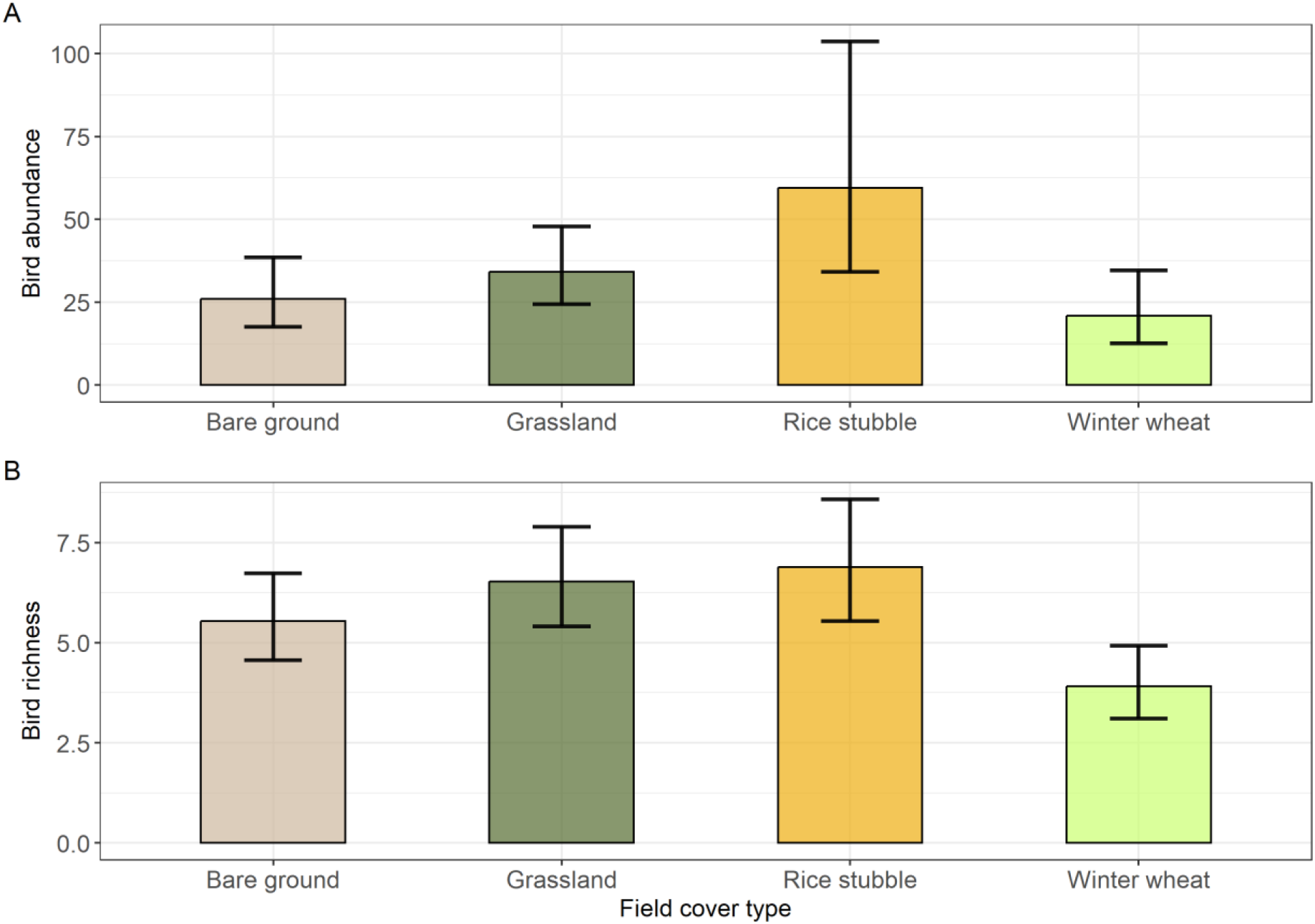
Abundance (A) and richness (B) of wintering birds in the agricultural fields of the Camargue depending on different field cover types. Vertical bars delimit the 95% confidence intervals.

Species richness was also explained by field cover type (Table 1) with a significantly lower richness in winter wheat fields compared to the other field cover types (*p* < 0.01 for all pairwise comparisons between winter wheat and other cover types; Fig. 2b). Field cover diversity and landscape feature diversity were retained in the set of models within ΔAICc=2 but were discarded when proceeding to model averaging, which suggests low support to these effects (ESM Table S2).

### Species-specific response

The effect of stubble field on species abundance varied depending on species identity (Table 2; Fig. 3).

**Figure 3.**
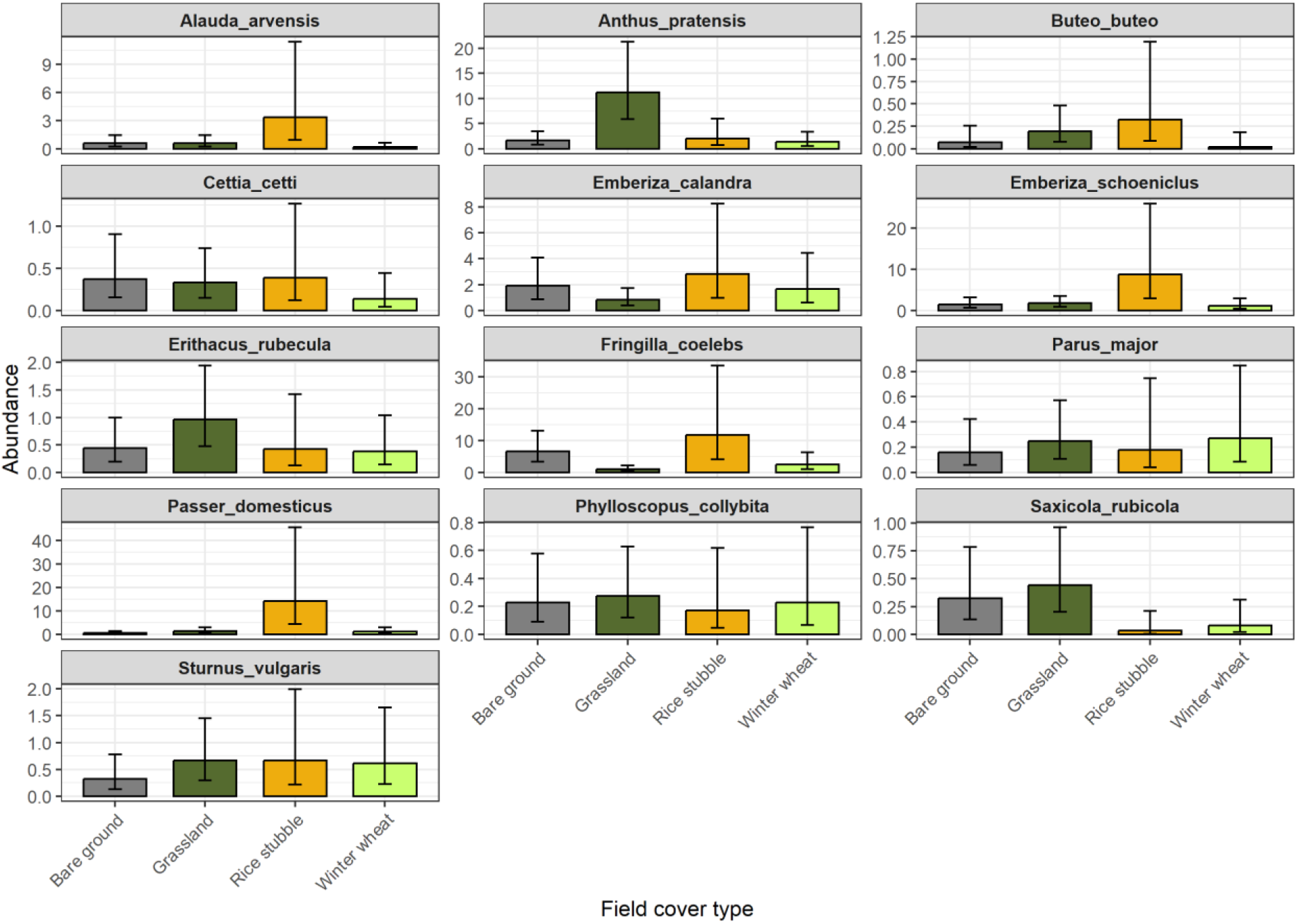
Abundance of wintering bird species in the crop fields of the Camargue depending on the different winter crop covers for each species.

**Table 2.** Model name, effects, number of parameters, AICc and ΔAICc for models assessing the effects of landscape features at the field level (10m buffer) and at the landscape level (500m buffer) on bird abundance. In all models, effects are in interaction with species as fixed effect.

| **Model** | **Field cover effects** | **Landscape feature effects** | **k** | **AICc** | **ΔAIC** |
|---|---|---|---|---|---|
| **M1** | Field cover type +Stubble area | Hedgerow10+Hedgerow500+Grass_strip10+Reed_strip10+Reed_strip500 | 135 | 5998.8 | 0 |
| **M2** | Field cover type +Stubble area | Hedgerow10+Hedgerow500+Grass_strip500+Reed_strip10+Reed_strip500 | 135 | 6004.9 | 6.1 |
| **M3** | Field cover type +Stubble area | Hedgerow10+Hedgerow500+Grass_strip500+Reed_strip500 | 122 | 6008.0 | 9.2 |
| **M4** | Field cover type +Stubble area | Hedgerow10+Hedgerow500+Grass_strip10+Grass_strip500+Reed_strip500 | 135 | 6009.4 | 10.6 |
| **M6** | Field cover type + Stubble area | Hedgerow10+Hedgerow500+Grass_strip10+Grass_strip500 | 122 | 6011.0 | 12.2 |
| **M5** | Field cover type + Stubble area | Hedgerow10+Hedgerow500+Grass_strip10+Grass_strip500+Reed_strip10+ Reed_strip500 | 148 | 6012.0 | 13.2 |
| **M7** | Field cover type + Stubble area | Hedgerow10+Grass_strip10+Grass_strip500+Reed_strip10+Reed_strip500 | 135 | 6013.5 | 14.7 |
| **M8** | Field cover type + Stubble area | Hedgerow10+Hedgerow500+Grass_strip10+Grass_strip500+Reed_strip10 | 135 | 6018.1 | 19.2 |
| **M9** | Field cover type + Stubble area | Hedgerow500+Grass_strip10+Grass_strip500+Reed_strip10+ Reed_strip500 | 135 | 6029.0 | 30.2 |

Pairwise post-hoc contrasts showed that Eurasian skylarks and Reed buntings were more abundant in rice stubble than in winter wheat fields (respectively *p* = 0.015 and *p* = 0.03). Common chaffinchs were significantly more abundant in stubble fields and bare ground than in grasslands (*p* < 0.002) while House sparrows were significantly more abundant in stubble fields than in all other cover types (*p* < 0.002; Fig. 3). Meadow pipits were significantly more abundant in grassland fields than in the other types of field cover (*p* < 0.04). For all other species, there were no significant difference in abundance among field cover types.

Rice stubble area over the landscape had a significant effect on species abundance only for one species, decreasing the abundance of Skylarks (β = -1.17x10^-3^ ± 3.33x10^-6^, corresponding to 1 fewer bird per 15 ha of extra rice stubble; Fig. 4).

**Figure. 4.**
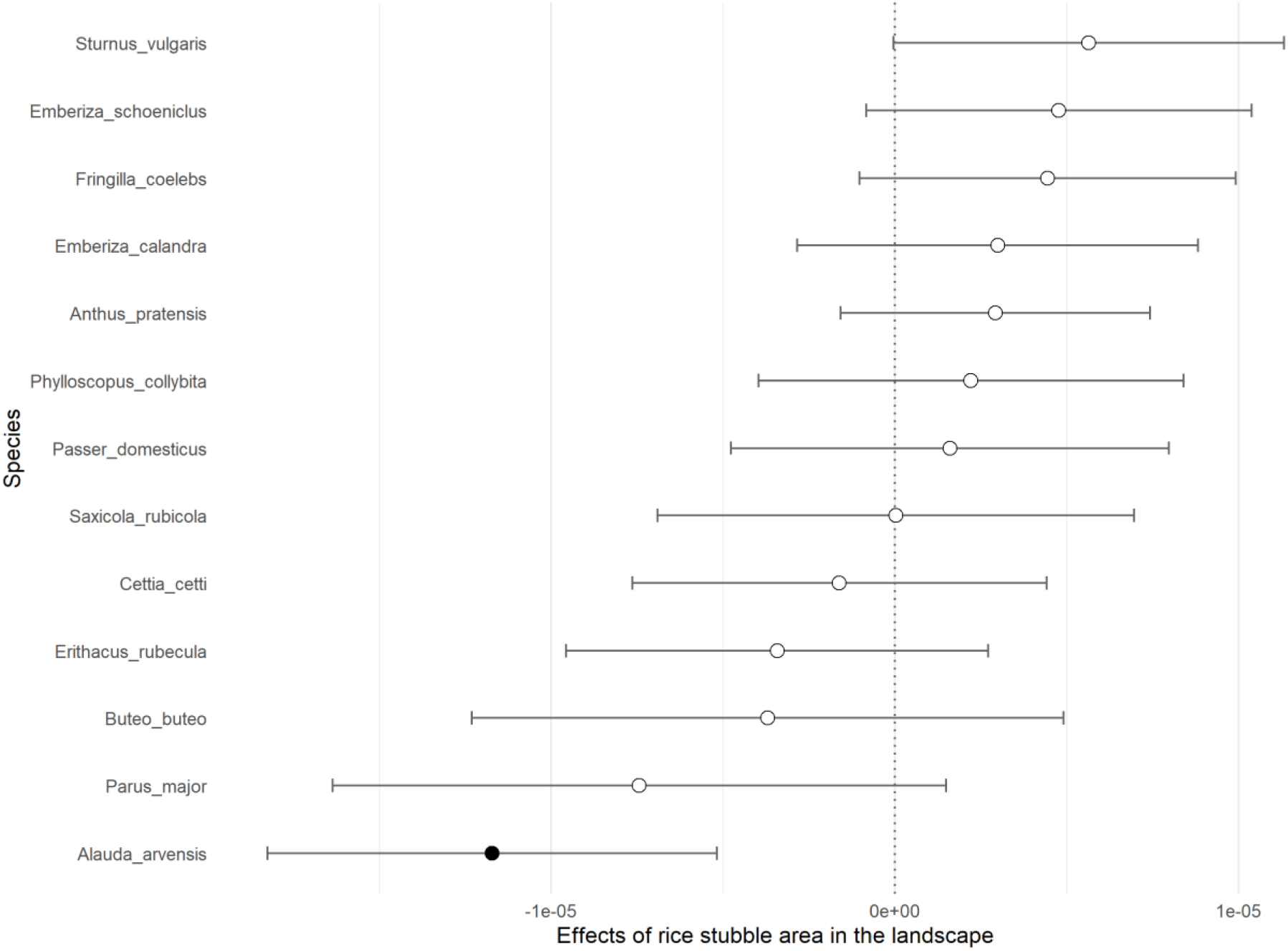
Species-specific slopes (± 95%CI) for the relationship between abundance and stubble area at the landscape scale (500m buffer) for wintering birds in the Camargue. Black circles are for slopes significantly different from zero.

At the field level there was an effect of hedgerows, reed strips and grass strips area in interaction with species (Table 2; Fig. 5). Species which benefited from increasing hedgerow area at field level were Common starling, Great tit, Common chiffchaff, and Common chaffinch (from 1 more bird per 0.2 ha extra hedgerow for Common starling to 1 more bird per 0.4 ha extra surface area of hedgerow for Common chaffinch; Fig. 5a). None of the species had a negative relationship of abundance with hedgerows area (Fig. 5a). Reed strips area along the field edge had a positive effect on Common starlings and European stonechats (from 0.3 to 0.5 ha of extra surface area for one more bird, respectively) while having a negative effect on Corn buntings and Common sparrow (from 0.3 to 0.7 ha of extra surface for one bird less, respectively; Fig. 5c). Grass strips area along the field edge had a positive effect on Eurasian skylarks and Reed buntings (0.25 ha and 0.5 ha of extra surface area for one more bird, respectively) and a negative effect on Common chaffinches (with 0.4 ha of extra surface for one bird less; Fig. 5e).

**Figure 5.**
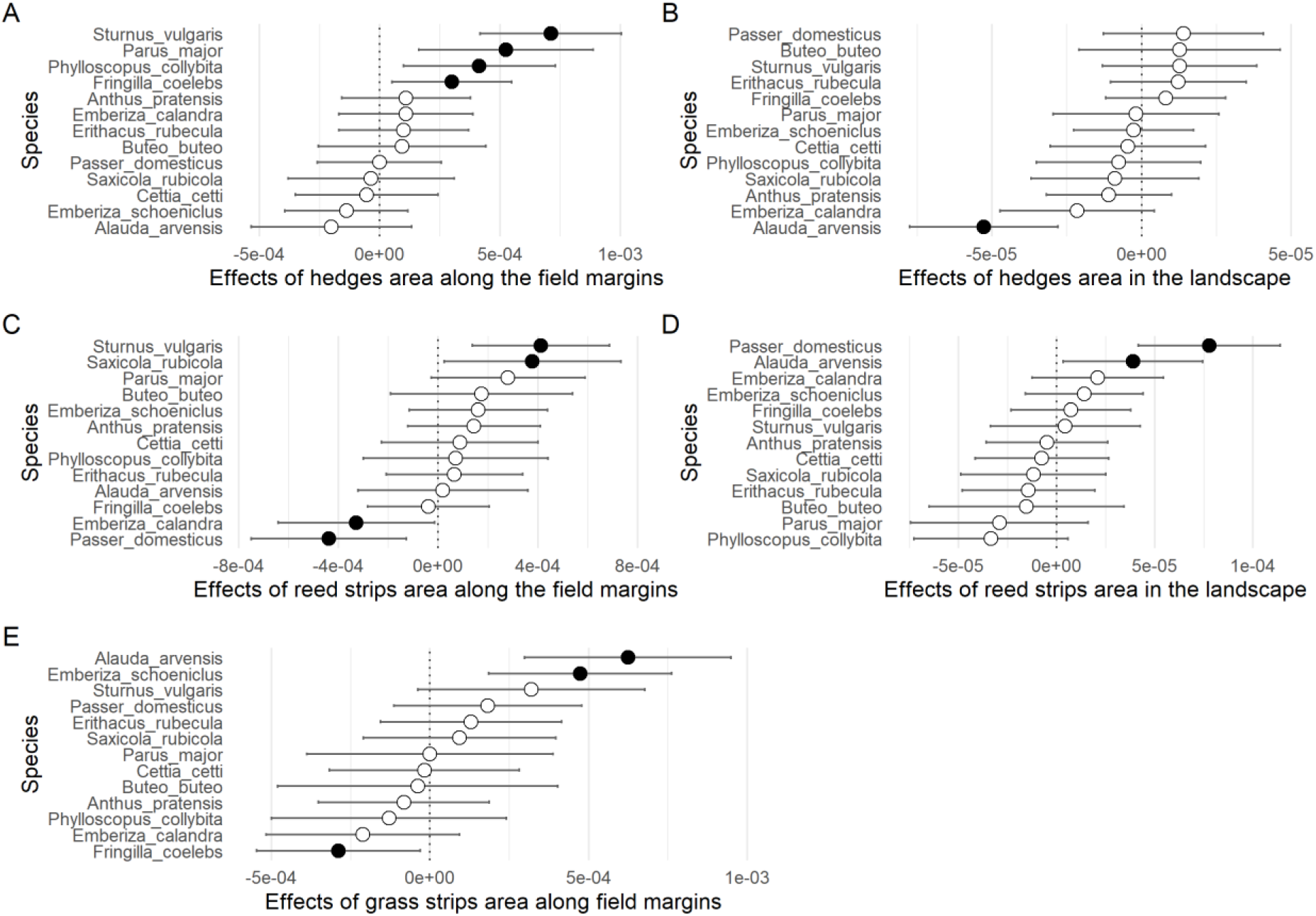
Species-specific slopes (± 95%CI) for the relationship between abundance of wintering birds and A) hedgerow area along the field margins (10m buffer), B) hedgerow area in the landscape (500m buffer), C) reed strips area along the field margins, D) reed strips area in the landscape and E) grass strips area along the field margins. Black circles are for slopes significantly different from zero.

At the landscape level, the hedgerow area was only significantly negative for Eurasian skylarks (with one bird less per 2 ha of extra surface area of hedgerow; Fig. 5b). Finally, the effect of reed strips area was retained at the landscape level with a positive effect for House sparrows and Eurasian skylark (7 ha and 5 ha of extra surface area of reed for one more bird, respectively, Fig. 5d).

## Discussion

Our study shows that, in the Camargue, rice stubble fields accommodate higher abundance of wintering birds than every other field cover type in line with our first prediction. Moreover, our study shows that this pattern is driven by a few gregarious and granivorous wintering species. Second, we found that landscape feature diversity increases wintering bird abundance but has no effect on species richness at the community level thus partially supporting the complementation hypothesis and our third prediction. Together, these results highlight the complementary roles of field- and landscape-level management for wintering bird communities in the Camargue. Finally, we emphasize species-specific responses at both the field and landscape level, which can be used to develop targeted management recommendations.

### Field cover type

Our results show that the presence of rice stubble fields has a positive effect on wintering bird abundance. In the Camargue, Pernollet et al. (2017) found that stubble rice fields contain both waste rice seeds and seeds of weeds. An average of 349.9 ± 57.6 kg/ha of rice seeds is left in the fields after harvest, while weed seed densities average 141.6 ± 21.3 kg/ha with 3 species representing 82% of the mean weed seed density (47% of red rice, 25% of barnyard grass, and 10% cosmopolitan bulrush). Contrary to other regions of rice production where rice fields are flooded in winter, this practice is very reduced in the Camargue. While the limited winter flooding of rice fields may reduce their carrying capacity for wintering waterbirds (Pernollet et al. 2015), dry rice stubbles nevertheless provide valuable foraging habitat for several wintering passerines, particularly granivorous species. The leftover rice grains and weed seed, are available for wintering birds only if the usual post harvesting ploughing, which normally takes place in October, is postponed until February or March due to autumn heavy rainfalls.

The positive effect of stubble for granivorous wintering passerines is consistent with previous studies (Robinson and Sutherland 1997; Maeda 2001; Kasprzykowski and Goławski 2012). This positive effect is also consistent with the fact that stubble offers more food resources in terms of grain leftover than other types of field cover (Moorcroft et al. 2002). The abondance of waste rice grain may also have driven the recent increase in wintering Common cranes *Grus grus* in the Camargue (Salvi 2023). In Canada, the conversion of natural marshes into corn fields partly explains the increase of the Greater snow goose *Anser caerulescens caerulescens* populations observed since 1970. There, leftover grains in stubble cornfields provide abundant food resource to wintering geese (Gauthier et al. 2005). Increased carrying capacity offered by stubble fields in winter may play a role in population dynamics beyond the winter survival of these species as winter resource availability was shown to be one of the main drivers of the breeding populations of farmland birds (Gillings et al. 2005; Siriwardena et al. 2008b).

Yet, the area of stubble available for birds in winter is also decreasing in the Camargue due to the shift from spring-sown to autumn-sown dry cereals (such as wheat or barley), which offer higher yields (Beres et al. 2010) but are associated with earlier stubble ploughing (Newton 2004). Elsewhere it has been shown that food availability in stubble fields is negatively related to herbicides used during the growing season and improved harvest tools that spill fewer seeds (Stoate et al. 2009; Goded et al. 2018).

Meadow pipit was the only species showing a clear preference for grasslands. This is consistent with their broad winter diet, which includes both invertebrates and seeds, and with previous observations of preferential use of alfalfa and grassland fields (Powolny et al. 2018). Whether this preference reflects differences in seed availability, invertebrate abundance or vegetation structure cannot be disentangled from our data and would require further investigation.

While landscape feature diversity increased bird abundance its effect on richness was not supported. This contrast with other studies which found higher richness with increasing landscape feature diversity (Li et al. 2020) or natural vegetation patches (Morales et al 2015). In our context the presence of stubble fields may be so important as a food resource that it attracts large bird community independently of the landscape context. Alternatively, contrasting landscape features requirements among species with different life history traits may result in similar richness across a gradient of landscape features even if species identity differ. A species specific approach is required to disentangle this.

### Species-specific response to landscape features at different levels

Consistent with the complementation hypothesis, we found that the area of field margins with dense vegetation, i.e. hedgerow and reed strips, had a positive effect on the abundance of several species. As expected, species favored by hedgerows at the field level are generalist species of forest edges (i.e. Common starling, Great tit, Common chiffchaff and Common Chaffinch; Fig. 5a). This results is in line with other studies conducted in different agricultural contexts, e.g. annual crops in central Chile during the austral winter (Muñoz-Sáez et al. 2017) and in orchards in Switzerland (Rime et al. 2020). During winter, hedgerows offer shelter against predators but also food resources such as insects using trees and bushes for overwintering (Maudsley et al. 2002; Gallé et al. 2018). The presence of fruit trees such as blackthorn *Prunus spinosa* in hedgerows of the Camargue could contribute to this positive effect.

Hedgerows have been shown to favor some species but deter other species and this may depend on the context. Indeed, a study conducted in Spanish vineyards found a negative effect of hedgerows on insectivorous birds (Rollan et al. 2019) while a positive effect of hedgerows was observed in Switzerland orchard landscapes (Rime et al. 2020).

Similarly, the dense structure of reed strips could be used as shelter for several species. Our results show a positive effect of this landscape feature at the field level for Common starling and European stonechat (Fig. 5c). Common starlings are known to roost in reedbeds in the Camargue so that their presence may be related to this behavior. European stonechat may prefer the low level of reed strips to hedgerows for hunting insects that they may also found along irrigation and drainage canals. Indeed, reed strips could also provide supplementary food as they host many invertebrates (spiders, ground beetles, and long-legged flies) at least during the breeding season (Schmidt et al. 2005; Decleer et al. 2015).

At the field level grass strips were found to favor Skylarks and Reed buntings, an expected results for two species of open habitat (Fig 5e). In Poland, Reed bunting diet was found to be mainly composed of seeds of *Amaranthus retroflexus* and *Chenopodium album*, two weeds that become more common due to herbicide resistance (Orłowski and Czarnecka 2007) and are also common in fallow and grass strips along field margins of the Camargue (Marnotte et al. 2006).

At the landscape level (Fig. 5b & 5d), only hedgerow and reed strips were retained in the best model. While the negative response of Skylarks to hedgerow was expected because of the affinity of skylarks with open areas, it is less clear why it respond positively to reed strips. It may simply result from the negative correlation (*r* = - 0.49; ESM Fig. S1) between both landscape features. The same process may explain the significant negative effect of stubble area at the landscape scale for this species as this parameter also tends to be negatively correlated with hedgerow area (*r* = - 0.40; ESM Fig. S1)

Overall, our results highlight the importance of maintaining a several types of landscape features in the landscape if one aims at preserving a wide community of species, as it was already found for breeding birds in the same Camargue landscape (Mallet et al. 2023). The fact that species respond more strongly to landscape features in the immediate vicinity of the field than to those in the landscape suggests that these different scales should be taken into account in future studies. Whilst the importance of the landscape level is undeniable during the breeding season (Batáry et al. 2011; Sirami et al. 2019), this factor may be less significant outside this period, when individuals are less attached to their territory.

### Management recommendations

Developing recommendations for biodiversity friendly practices to farmer is necessarily a highly context dependent process, which makes this first benchmark study in a wetland agricultural system so important. This work could contribute to design new agri-environmental schemes that help increasing both the abundance and the diversity of wintering birds in rice production areas. Our results suggest that agri-environmental schemes should not focus on a single landscape feature, but rather aim to maintain complementary resources at both field and landscape levels. Delaying stubble ploughing could maintain important winter food resources for granivorous birds, while preserving a mosaic of grasslands, winter crops and different landscape features would accommodate species with contrasting habitat requirements. Therefore, supporting schemes that are built at such a landscape-level approach may therefore simultaneously support high bird abundance and community diversity.

For instance, new agri-environmental schemes could compensate farmers for delaying the ploughing of rice stubble, thereby maintaining food resources throughout winter. While this kind of agri-environmental scheme has been experimented in the UK with good success (Baker et al. 2012), it is not part of the portfolio of measures available to rice farmers in the Camargue. This subsidy would help them overcome the financial risk of delaying crop tillage operations, since it may be more difficult to achieve crop tillage in early spring when rainfall are frequent, hampering soil preparation. Delaying ploughing would also help to minimize the damage caused by Common cranes to autumn-sown crop seedlings by concentrating the birds on rice stubble. Ensuring that beyond conserving rice stubble, a diversity of weed species remain available (through a further support to organic practices) is also key to expect positive results for wintering birds (Hancock and Wilson 2003). Coupling such support to already existing schemes supporting hedgerow maintenance or to new ones supporting grass strip enlargement could help reverse wintering bird passerine decline like it was done for the Cirl bunting *Emberiza cirlus* in the UK (Peach et al. 2001).

Because, organic rice farmers try to diversify their crop rotations, planting winter seed-rich winter bird crops could also be encouraged as it was experimented in England with Kale or Quinoa (Henderson et al. 2004), two crops that may be cultivated in the Camargue.

## Acknowledgements

We are grateful to Biosud, one of the main collector of organic rice in the Camargue. Biosud favoured the linked between our research team and the farmers who accepted to participate in this study. We also thank all farm landowners who graciously permitted us to work in their fields.

---

**Figure S1.**
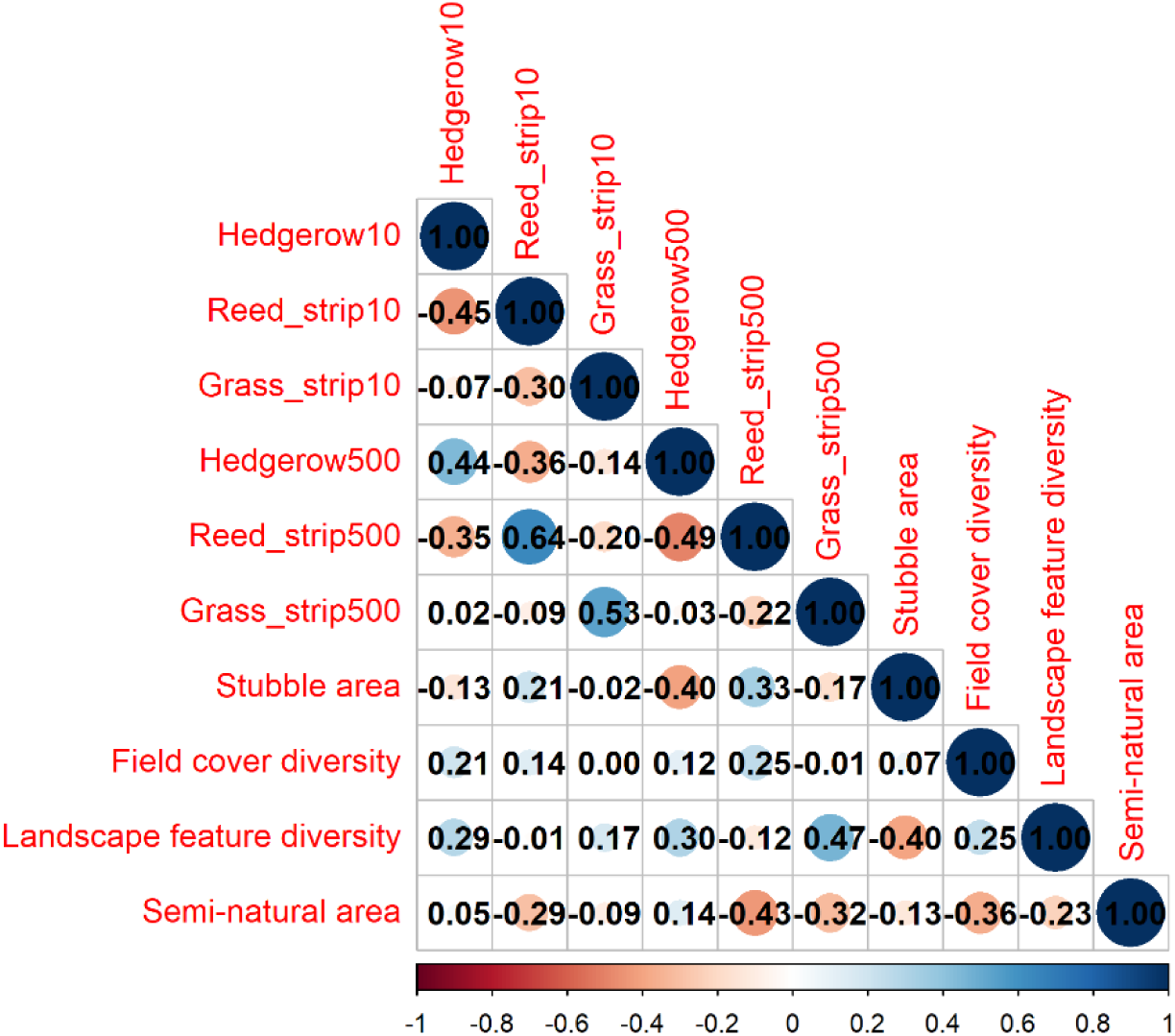
Correlation table among explanatory variables.

**Table S1.** List of the species detected during two visits in winter 2021 and 2022 in 86 fields of the Camargue (South of France).

| **Species** | **min** | **mean** | **max** | **occurrence** | **frequence** |
|---|---|---|---|---|---|
| **Anthus_pratensis** | 0 | 5.5 | 111 | 95 | 55.2 |
| **Fringilla_coelebs** | 0 | 9.3 | 255 | 90 | 52.3 |
| **Emberiza_schoeniclus** | 0 | 6.0 | 146 | 83 | 48.3 |
| **Erithacus_rubecula** | 0 | 0.7 | 5 | 78 | 45.3 |
| **Cettia_cetti** | 0 | 0.3 | 2 | 50 | 29.1 |
| **Passer_domesticus** | 0 | 7.7 | 190 | 50 | 29.1 |
| **Alauda_arvensis** | 0 | 3.3 | 120 | 40 | 23.3 |
| **Emberiza_calandra** | 0 | 4.1 | 120 | 37 | 21.5 |
| **Saxicola_rubicola** | 0 | 0.3 | 4 | 36 | 20.9 |
| **Phylloscopus_collybita** | 0 | 0.6 | 27 | 32 | 18.6 |
| **Sturnus_vulgaris** | 0 | 2.5 | 170 | 31 | 18.0 |
| **Parus_major** | 0 | 0.6 | 36 | 30 | 17.4 |
| **Buteo_buteo** | 0 | 0.2 | 3 | 26 | 15.1 |
| **Sylvia_atricapilla** | 0 | 0.3 | 5 | 24 | 14.0 |
| **Curruca_melanocephala** | 0 | 0.2 | 2 | 23 | 13.4 |
| **Carduelis_carduelis** | 0 | 0.5 | 19 | 22 | 12.8 |
| **Cisticola_juncidis** | 0 | 0.2 | 5 | 21 | 12.2 |
| **Troglodytes_troglodytes** | 0 | 0.1 | 3 | 18 | 10.5 |
| **Turdus_merula** | 0 | 0.2 | 6 | 17 | 9.9 |
| **Motacilla_alba** | 0 | 0.4 | 25 | 16 | 9.3 |
| **Phoenicurus_ochruros** | 0 | 0.1 | 4 | 16 | 9.3 |
| **Cyanistes_caeruleus** | 0 | 0.1 | 2 | 14 | 8.1 |
| **Galerida_cristata** | 0 | 0.1 | 3 | 14 | 8.1 |
| **Ardea_cinerea** | 0 | 0.1 | 2 | 12 | 7.0 |
| **Turdus_philomelos** | 0 | 0.2 | 10 | 11 | 6.4 |
| **Streptopelia_decaocto** | 0 | 0.5 | 70 | 9 | 5.2 |
| **Chloris_chloris** | 0 | 0.1 | 3 | 8 | 4.7 |
| **Corvus_corone** | 0 | 0.2 | 22 | 8 | 4.7 |
| **Falco_tinnunculus** | 0 | 0.0 | 1 | 8 | 4.7 |
| **Passer_montanus** | 0 | 0.3 | 25 | 8 | 4.7 |
| **Aegithalos_caudatus** | 0 | 0.2 | 10 | 6 | 3.5 |
| **Dryobates_minor** | 0 | 0.0 | 2 | 6 | 3.5 |
| **Fringilla_montifringilla** | 0 | 0.0 | 3 | 6 | 3.5 |
| **Pica_pica** | 0 | 0.1 | 14 | 6 | 3.5 |
| **Accipiter_nisus** | 0 | 0.0 | 1 | 5 | 2.9 |
| **Bubulcus_ibis** | 0 | 0.1 | 12 | 5 | 2.9 |
| **Grus_grus** | 0 | 0.3 | 33 | 5 | 2.9 |
| **Regulus_ignicapilla** | 0 | 0.0 | 2 | 5 | 2.9 |
| **Casmerodius_albus** | 0 | 0.2 | 26 | 4 | 2.3 |
| **Gallinago_gallinago** | 0 | 0.1 | 6 | 4 | 2.3 |
| **Phasianus_colchicus** | 0 | 0.1 | 8 | 4 | 2.3 |
| **Columba_palumbus** | 0 | 0.4 | 70 | 3 | 1.7 |
| **Corvus_monedula** | 0 | 0.4 | 54 | 3 | 1.7 |
| **Emberiza_cirlus** | 0 | 0.1 | 12 | 3 | 1.7 |
| **Picus_viridis** | 0 | 0.0 | 1 | 3 | 1.7 |
| **Prunella_modularis** | 0 | 0.0 | 1 | 3 | 1.7 |
| **Regulus_regulus** | 0 | 0.0 | 2 | 3 | 1.7 |
| **Alcedo_atthis** | 0 | 0.0 | 1 | 2 | 1.2 |
| **Dendrocopos_major** | 0 | 0.0 | 1 | 2 | 1.2 |
| **Larus_michahellis** | 0 | 0.0 | 5 | 2 | 1.2 |
| **Serinus_serinus** | 0 | 0.0 | 1 | 2 | 1.2 |
| **Turdus_pilaris** | 0 | 0.0 | 2 | 2 | 1.2 |
| **Upupa_epops** | 0 | 0.0 | 3 | 2 | 1.2 |
| **Vanellus_vanellus** | 0 | 0.0 | 6 | 2 | 1.2 |
| **Ciconia_ciconia** | 0 | 0.0 | 2 | 1 | 0.6 |
| **Circus_aeruginosus** | 0 | 0.0 | 2 | 1 | 0.6 |
| **Circus_cyaneus** | 0 | 0.0 | 1 | 1 | 0.6 |
| **Columba_livia** | 0 | 0.1 | 19 | 1 | 0.6 |
| **Coturnix_coturnix** | 0 | 0.0 | 2 | 1 | 0.6 |
| **Elanus_caeruleus** | 0 | 0.0 | 2 | 1 | 0.6 |
| **Emberiza_citrinella** | 0 | 0.0 | 2 | 1 | 0.6 |
| **Luscinia_svecica** | 0 | 0.0 | 1 | 1 | 0.6 |
| **Motacilla_cinerea** | 0 | 0.0 | 1 | 1 | 0.6 |
| **Rallus_aquaticus** | 0 | 0.0 | 1 | 1 | 0.6 |
| **Tringa_ochropus** | 0 | 0.0 | 1 | 1 | 0.6 |

**Table S2.** Estimate ± SE and 95% confidence interval after model averaging of the models within 2 points of AICc for total bird abundance and richness.

| **Response variable** | **Effect** | **Estimate ± SE** | **95% CI** |
|---|---|---|---|
| **Total bird abundance** | Intercept | 3.414 ± 0.192 | [3.034 ; 3.794] |
|  | Field cover type_grassland | 0.089 ± 0.202 | [-0.329 ; 0.673] |
|  | Field cover type_stubble | **0.371 ± 0.439** | [0.007 ; 1.418] |
|  | Field cover type_wheat | -0.120 ± 0.234 | [-0.789 ; 0.328] |
|  | Field cover diversity | 0.052 ± 0.095 | [-0.096 ; 0.344] |
|  | Landscape feature diversity | **0.198 ± 0.136** | [0.004 ; 0.460] |
|  | Stubble area | **0.408 ± 0.146** | [0.120 ; 0.696] |
| **Richness** | Intercept | 1.713 ± 0.097 | [1.520 ; 1.906] |
|  | Field cover type_grassland | 0.164 ± 0.094 | [-0.022 ; 0.350] |
|  | **Field cover type_stubble** | **0.216 ± 0.107** | **[0.003 ; 0.429]** |
|  | **Field cover type_wheat** | **-0.340 ± 0.114** | **[-0.566 ; -0.113]** |
|  | Field cover diversity | 0.027 ± 0.039 | [-0.011 ; 0.136] |
|  | Landscape feature diversity | 0.046 ± 0.042 | [-0.036 ; 0.129] |

## References

1. Baker DJ, Freeman SN, Grice PV, Siriwardena GM (2012) Landscape-scale responses of birds to agri-environment management: a test of the English Environmental Stewardship scheme. Journal of Applied Ecology 49:871–882. 10.1111/j.1365-2664.2012.02161.x

2. Barton K (2020) MuMIn: Multi-model inference. R package version 1.43.6

3. Batáry P, Fischer J, Báldi A, et al (2011) Does habitat heterogeneity increase farmland biodiversity? Frontiers in Ecol & Environ 9:152–153. 10.1890/11.WB.006

4. Benton TG, Bryant DM, Cole L, Crick HQP (2002) Linking agricultural practice to insect and bird populations: a historical study over three decades. Journal of Applied Ecology 39:673–687. 10.1046/j.1365-2664.2002.00745.x

5. Beres BL, Harker KN, Clayton GW, et al (2010) Weed-competitive ability of spring and winter cereals in the Northern Great Plains. Weed Technology 24:108–116. 10.1614/wt-d-09-00036.1

6. Burnham KP, Anderson DR (2002) Model selection and multimodel inference, A practical information-theoretic approach. 2nd ed Springer-Verlag, New-York

7. Chamberlain DE, Wilson AM, Browne SJ, Vickery JA (1999) Effects of habitat type and management on the abundance of skylarks in the breeding season. Journal of Applied Ecology 36:856–870. 10.1046/j.1365-2664.1999.00453.x

8. Decleer K, Maes D, Van Calster H, et al (2015) Importance of core and linear marsh elements for wetland arthropod diversity in an agricultural landscape. Insect Conservation and Diversity 8:289–301. 10.1111/icad.12110

9. Donald PF, Evans AD (1994) Habitat selection by Corn Buntings *Miliaria calandra* in winter. Bird Study 41:199–210. 10.1080/00063659409477220

10. Dunning JB, Danielson BJ, Pulliam HR (1992) Ecological processes that effect populations in complex landscapes. Oikos 65:169–175. 10.2307/3544901

11. Fraixedas S, Galewski T, Ribeiro-Lopes S, et al (2019) Estimating biodiversity changes in the Camargue wetlands: An expert knowledge approach. PLoS ONE 14:e0224235. 10.1371/journal.pone.0224235

12. Gallé R, Császár P, Makra T, et al (2018) Small-scale agricultural landscapes promote spider and ground beetle densities by offering suitable overwintering sites. Landscape Ecology 33:1435–1446. 10.1007/s10980-018-0677-1

13. Gauthier G, Giroux JF, Reed A, et al (2005) Interactions between land use, habitat use, and population increase in greater snow geese: What are the consequences for natural wetlands? Global Change Biology 11:856–868. 10.1111/j.1365-2486.2005.00944.x

14. Geiger F, De Snoo GR, Berendse F, et al (2010) Landscape composition influences farm management effects on farmland birds in winter: A pan-European approach. Agriculture, Ecosystems & Environment 139:571–577. 10.1016/j.agee.2010.09.018

15. Gillings S, Newson SE, Noble DG, Vickery JA (2005) Winter availability of cereal stubbles attracts declining farmland birds and positively influences breeding population trends. Proceedings of the Royal Society B: Biological Sciences 272:733–739. 10.1098/rspb.2004.3010

16. Goded S, Ekroos J, Domínguez J, et al (2018) Effects of organic farming on bird diversity in North-West Spain. Agriculture, Ecosystems and Environment 257:60–67. 10.1016/j.agee.2018.01.020

17. Hancock MH, Wilson JD (2003) Winter habitat associations of seed-eating passerines on Scottish farmland. Bird Study 50:116–130. 10.1080/00063650309461303

18. Haviland DR, Caprile JL, Rill SM, et al (2016) Phenology of spotted wing drosophila in the San Joaquin Valley varies by season, crop and nearby vegetation. California Agriculture 70:24–31. 10.3733/ca.v070n01p24

19. Henderson IG, Vickery JA, Carter N (2004) The use of winter bird crops by farmland birds in lowland England. Biological Conservation 118:21–32. 10.1016/j.biocon.2003.06.003

20. Issa N, Muller Y (2015) Atlas des oiseaux nicheurs de France métropolitaine

21. IUCN (2024) Agriculture and conservation. Gland

22. Jiguet F (2014) Suivi hivernal des oiseaux communs

23. Kasprzykowski Z, Goławski A (2012) Habitat preferences of granivorous passerine birds wintering on farmland in eastern Poland. Bird Study 59:52–57. 10.1080/00063657.2011.643292

24. Katayama N, Osawa T, Amano T, Kusumoto Y (2015) Are both agricultural intensification and farmland abandonment threats to biodiversity? A test with bird communities in paddy-dominated landscapes. Agriculture, Ecosystems and Environment 214:21–30. 10.1016/j.agee.2015.08.014

25. Kristensen K, McGillycuddy M (2022) Covariance structures with glmmTMB. https://cran.r-project.org/web/packages/glmmTMB/vignettes/covstruct.html. Accessed 4 Aug 2022

26. Lenth VL, Buerkner P, Herve M, et al (2022) Package ‘emmeans.’ 94

27. Li D, Lee M, Xiao W, et al (2020) Noncrop features and heterogeneity mediate overwintering bird diversity in agricultural landscapes of southwest China. Ecology and Evolution 10:5815–5828. 10.1002/ece3.6319

28. Maeda T (2001) Patterns of bird abundance and habitat use in rice fields of the Kanto Plain, central Japan. Ecological Research 16:569–585. 10.1046/j.1440-1703.2001.00418.x

29. Magnusson A, Skaug H, Nielsen A, et al (2020) Generalized linear mixed models using template model builder. Package glmmTMB. Version 1.0.2.1. In: R topics documented. https://cran.r-project.org/package=glmmTMB. Accessed 19 Oct 2021

30. Mallet P, Bechet A, Sirami C, et al (2023) Field margins as substitute habitat for the conservation of birds in agricultural wetlands. Peer Community Journal 3:. 10.24072/pcjournal.299

31. Marcon E, Hérault B (2015) Entropart: An R package to measure and partition diversity. Journal of Statistical Software 67:. 10.18637/jss.v067.i08

32. Marja R, Herzon I (2012) The importance of drainage ditches for farmland birds in agricultural landscapes in the Baltic countries: Does field type matter? Ornis Fennica 89:170–181

33. Marnotte P, Carrara A, Dominati E, Girardot F (2006) Plantes des rizières de Camargue, Quae

34. Maudsley M, Seeley B, Lewis O (2002) Spatial distribution patterns of predatory arthropods within an English hedgerow in early winter in relation to habitat variables. Agriculture, Ecosystems and Environment 89:77–89. 10.1016/S0167-8809(01)00320-6

35. Moorcroft D, Whittingham MJ, Bradbury RB, Wilson JD (2002) The selection of stubble fields by wintering granivorous birds reflects vegetation cover and food abundance. Journal of Applied Ecology 39:535–547. 10.1046/j.1365-2664.2002.00730.x

36. Morales MB, Oñate JJ, Guerrero I, Meléndez L (2015) Influence of landscape and field-level agricultural management on a mediterranean farmland winter bird community. Ardeola 62:49–65. 10.13157/arla.62.1.2015.49

37. Moreau J, Monceau K, Gonnet G, et al (2022) Organic farming positively affects the vitality of passerine birds in agricultural landscapes. Agriculture, Ecosystems and Environment 336:108034. 10.1016/j.agee.2022.108034

38. Mouret J-C (2018) Les exigences de la conduite d’une rizière en Camargue. In: Cardère (ed) Le riz et la Camargue. Vers des agrosystèmes durables, Educagri. pp 109–124

39. Muñoz-Sáez A, Perez-Quezada JF, Estades CF (2017) Agricultural landscapes as habitat for birds in central Chile. Revista Chilena de Historia Natural 90:. 10.1186/s40693-017-0067-0

40. Newton I (2017) Farming and birds. HarperCollins UK, London

41. Newton I (2004) The recent declines of farmland bird populations in Britain: An appraisal of causal factors and conservation actions. Ibis 146:579–600. 10.1111/j.1474-919X.2004.00375.x

42. Orłowski G, Czarnecka J (2007) Winter diet of reed bunting Emberiza schoeniclus in fallow and stubble fields. Agriculture, Ecosystems & Environment 118:244–248. 10.1016/j.agee.2006.05.026

43. Pasher J, Mitchell SW, King DJ, et al (2013) Optimizing landscape selection for estimating relative effects of landscape variables on ecological responses. Landscape Ecology 28:371–383. 10.1007/s10980-013-9852-6

44. Peach WJ, Lovett LJ, Wotton SR, Jeffs C (2001) Countryside stewardship delivers cirl buntings (Emberiza cirlus) in Devon, UK. Biological Conservation 101:361–373. 10.1016/S0006-3207(01)00083-0

45. Peach WJ, Siriwardena GM, Gregory RD (1999) Long-term changes in over-winter survival rates explain the decline of reed buntings *Emberiza schoeniclus* in Britain. Journal of Applied Ecology 36:798–811. 10.1046/j.1365-2664.1999.00445.x

46. PECBMS (2025) Species trends. https://pecbms.info/trends-and-indicators/species-trends/sort/taxonomy/. Accessed 5 Jan 2025

47. PNRC (2016) L’occupation du sol en Camargue, état 2016 et son évolution depuis 2001. http://geo.pnrpaca.org/geoservices/decouvrir-les-parcs/des-parcs-et-des-projets/occupation-du-sol-en-camargue-etat-2016-et-son-evolution-depuis-2001/. Accessed 9 Apr 2020

48. R Core Team (2017) R: A Language and Environment for Statistical Computing

49. Rime Y, Luisier C, Arlettaz R, Jacot A (2020) Landscape heterogeneity and management practices drive habitat preferences of wintering and breeding birds in intensively-managed fruit-tree plantations. Agriculture, Ecosystems and Environment 295:106890. 10.1016/j.agee.2020.106890

50. Robinson R, Sutherland WJ (1997) The feeding ecology of seed-eating birds on farmland in winter. In: The ecology and conservation of corn buntings. pp 162–169

51. Rodier R, Boutron O, Béchet A, Therond O (2025) Interdependencies among hydro-saline dynamics, economic activities, ecological processes, and biodiversity in a deltaic social-ecological system: insights from the Rhône delta (southern France). E&S 30:art11. 10.5751/ES-16122-300311

52. Rollan À, Hernández-Matías A, Real J (2019) Organic farming favours bird communities and their resilience to climate change in Mediterranean vineyards. Agriculture, Ecosystems and Environment 269:107–115. 10.1016/j.agee.2018.09.029

53. Salvi A (2023) Evolution of the status of the Common Crane Grus grus in France from the 1970s to the present day. Proceedings of X International Crane Conference, Tartu – Estonia 80–96

54. Schmidt MH, Lefebvre G, Poulin B, Tscharntke T (2005) Reed cutting affects arthropod communities, potentially reducing food for passerine birds. Biological Conservation 121:157–166. 10.1016/j.biocon.2004.03.032

55. Sirami C, Gross N, Baillod AB, et al (2019) Increasing crop heterogeneity enhances multitrophic diversity across agricultural regions. Proc Natl Acad Sci USA 116:16442–16447. 10.1073/pnas.1906419116

56. Siriwardena GM, Calbrade NA, Vickery JA (2008a) Farmland birds and late winter food: Does seed supply fail to meet demand? Ibis 150:585–595. 10.1111/j.1474-919X.2008.00828.x

57. Siriwardena GM, Calbrade NA, Vickery JA (2008b) Farmland birds and late winter food: Does seed supply fail to meet demand? Ibis 150:585–595. 10.1111/j.1474-919X.2008.00828.x

58. Stoate C, Báldi A, Beja P, et al (2009) Ecological impacts of early 21st century agricultural change in Europe - A review. Journal of Environmental Management 91:22–46. 10.1016/j.jenvman.2009.07.005

59. Vickery J, Carter N, Fuller RJ (2002) The potential value of managed cereal field margins as foraging habitats for farmland birds in the UK. Agriculture, Ecosystems and Environment 89:41–52. 10.1016/S0167-8809(01)00317-6

60. Vickery JA, Feber RE, Fuller RJ (2009) Arable field margins managed for biodiversity conservation: A review of food resource provision for farmland birds. Agriculture, Ecosystems & Environment 133:1–13. 10.1016/j.agee.2009.05.012

61. Whittingham MJ, Evans KL (2004) The effects of habitat structure on predation risk of birds in agricultural landscapes. Ibis 146:210–220. 10.1111/j.1474-919X.2004.00370.x

62. Wilson JD, Taylor R, Muirhead LB (1996) Field use by farmland birds in winter: An analysis of field type preferences using resampling methods. Bird Study 43:320–332. 10.1080/00063659609461025

63. Wuczyński A, Wuczyński M (2019) Birds wintering in heterogeneous farmland of Poland: Weather-dependent temporal changes in abundance and habitat associations. Acta Ornithologica 54:105–124. 10.3161/00016454AO2019.54.1.009

